# Disaggregation Regression and Multi-Model Evaluation for Predicting Dengue Risk in Africa

**DOI:** 10.1101/2025.06.17.660069

**Authors:** Jenicca Poongavanan, Tim C.D. Lucas, Gaspary Mwanyika, Moritz U.G. Kraemer, José Lourenço, Marcel Dunaiski, Tulio de Oliveira, Houriiyah Tegally, This is an initiative of the CLIMADE Consortium

## Abstract

Dengue risk mapping is essential for estimating disease burden, and informing targeted surveillance and control efforts. Current approaches to risk mapping vary widely in their methodology, data sources, output metrics and applications. Many existing approaches focus on predicting ecological suitability and produce high-resolution risk maps based on environmental conditions, yet high-resolution incidence maps remain scarce, leaving a critical gap in guiding precise, location-specific interventions. The prediction of disease incidence or transmission intensity remains relatively uncommon in disease ecology, largely due to data limitations, reporting biases, and the inherent complexity that arises from transmission dynamics. In this study, we applied disaggregation regression modelling to downscale aggregated dengue case data from 14 countries in Central and South America, generating fine-resolution incidence estimates that we subsequently projected onto the African continent. We then compared the resulting predictions from the incidence-based risk map with three widely used approaches: vector suitability index, dengue environmental suitability index, and mechanistic transmission potential (Index P). The disaggregation model achieved relatively strong predictive accuracy within the training region (mean correlation = 0.72) and showed partial alignment with reported burden across Africa (Spearman ρ = 0.33). Other risk maps exhibited similar or weaker correlations with reported cases in Africa, including ρ = 0.33 for dengue environmental suitability, ρ = 0.32 for transmission potential and ρ = 0.23 for Aedes aegypti suitability. Disaggregation regression offers a valuable tool for translating reported case data into spatially explicit estimates of burden, bridging the gap between ecological risk and epidemiological relevance. While spatial agreement was high in parts of coastal West Africa across the different risk map approaches, notable divergences highlight the distinct assumptions underlying each framework.

**Authors’ Summary:** Dengue is a mosquito-borne viral disease with expanding global impact. Accurately mapping dengue risk is essential for identifying areas of high transmission and targeting interventions effectively. Most current approaches to mapping dengue risk focus on environmental suitability for the virus or its mosquito vector, rather than estimating actual disease burden. In this study, we used an incidence-based approach; disaggregation regression, to estimate dengue cases at high spatial resolution using national and regional case data from Latin America. We then applied the model to Africa, where surveillance data are limited, and compared its predictions to three other common types of dengue risk maps. Our results showed that while all approaches provided some insight into geographic risk patterns, they often highlighted different areas as priorities. Our incidence-based model captured both where dengue might occur and how intense transmission may be, helping bridge the gap between environmental and ecological suitability of transmission and real-world disease burden. This approach can support more informed decision-making in areas with limited surveillance and guide targeted control efforts.

## Introduction

Dengue fever remains a significant global public health concern, with rising incidence and geographic expansion driven by urbanization, climate change, and human mobility [1,2]. Accurately mapping dengue risk is essential for informing surveillance, vector control strategies, and resource allocation in endemic and emerging regions. Over the past two decades, a range of approaches have been developed and applied to characterise spatio-temporal dengue risk, each leveraging different data sources and modeling frameworks [3,4].

These approaches can be broadly grouped into four categories based on their conceptual framework and data inputs: ecological niche models (ENMs) that use environmental covariates to predict the suitability of either vector or viral occurrence [5–8]; biologically-informed mechanistic models that estimate climate-driven transmission potential based on experimentally derived parameters like mosquito lifespan and incubation period [9,10]; incidence-based models that use reported case counts to estimate disease burden and transmission intensity [11–13]; and observational serological studies such as seroprevalence surveys and cohort analyses, which offer insights into population-level exposure and immunity. While the first two categories aim to predict areas of risk based on ecological or biological suitability, incidence-based models aim to quantify burden, providing estimates of where transmission is actually occurring. Despite differences in resolution, assumptions, and required inputs, they each provide a different and potentially complementary lens on where, when and how dengue risk emerges and evolves [14]. Lim et al. (2023) conducted a systematic review of the methodologies, covariates, and geographic extent of these different modeling approaches, offering a comprehensive synthesis of how *Aedes*-borne arbovirus transmission risk has been mapped to date [3]. A brief summary of their input data, pros, cons, and public health applications is provided in **Supp Table S1**.

These divergent methods reflect a broader complexity in dengue modeling, where multiple factors, including virus serotypes, mosquito vector ecology, human demographics, and socio-environmental conditions interact in nonlinear ways [15–17]. As highlighted by [14], comparing these models helps identify areas of agreement and divergence, assess the reliability of model predictions and clarify which representations of risk are most appropriate for specific decision-making contexts. This is particularly critical in settings like Africa, where dengue is historically underrecognized and underreported, yet evidence of endemic and epidemic transmission is growing [18–21].

In settings with limited data and high vulnerability, such comparative evaluations are critical to guide targeted interventions and resource allocation. For instance, ecological risk maps have been used to predict the likelihood of onward Zika virus transmission in Europe following case importations, thereby informing surveillance prioritization [22]. Transmission potential risk maps of dengue have also been leveraged to estimate the importation risk of dengue into non-endemic regions, supporting international preparedness and prioritization of ports of entry [23]. Moreover, effective dengue risk models ideally capture both spatial and temporal dynamics, incorporating climate, population, and socio-environmental factors across diverse geographies. However, the selection of appropriate covariates remains a core challenge, as models emphasise different mechanisms and relationships between the key aspects in transmission - host, vector and virus. These differing assumptions and structures often result in divergent risk predictions, raising important questions about model comparability and validity [14]. Rather than viewing such differences as contractions, they should be understood as a reflection of the varied conceptual frameworks and objectives each model brings. In this context, risk models should be seen as complementary tools, each offering distinct insights into dengue and more broadly, vector-borne disease transmission dynamics.

One key challenge is that ecological risk mapping for dengue estimates environmental suitability for transmission, but does not directly capture actual disease burden, as case numbers are also shaped by surveillance capacity, healthcare access and socio-demographic factors. Additionally, the relationship between the probability of dengue occurrence and the intensity of transmission may be non-linear, meaning areas predicted as highly suitable do not necessarily correspond to areas with high case counts. To explore how different risk mapping approaches characterize dengue risk in Africa, we compare three widely used ecological suitability models with an incidence-based model derived using a disaggregation regression approach. This statistical approach, which has shown promise for malaria burden estimation [24,25], downscales aggregated case data to finer spatial resolutions using covariate relationships. We train the model using detailed subnational case data from Central and South America and then apply it to Africa to generate high-resolution incidence estimates. It also allows for direct comparison with dengue risk maps and evaluation of how well a model trained in a data-rich setting performs in a data-limited context.

The dengue risk indicators used for comparison includes: (1) dengue suitability index built from dengue occurrence data [6], (2) vector suitability index representing environmental suitability for *Aedes aegypti*, the primary dengue vector [5]; while not a direct measure of dengue risk, it offers insight into the ecological potential for transmission, and (3) transmission potential map derived from mechanistic modelling of experimental mosquito-virus interactions (*Aedes Aegypti)* [26].

Beyond comparing modeling approaches, a key contribution of this study is the generation of high-resolution dengue incidence predictions for Africa, where such estimates are largely unavailable. By applying a disaggregation regression framework, we offer a practical tool that can support public health decision-making in settings where surveillance data are limited. At the same time, this work provides a critical look at how models developed in data-rich regions perform when applied to a different context, and whether more regionally tailored approaches may be needed. In doing so, we hope to contribute both a useful product and a broader reflection on how best to map and respond to dengue risk in under-surveilled parts of the world.

## Results

### Dengue Disaggregation Regression in South America, Model Performance and External Validation Across Independent Countries

To estimate fine-scale dengue burden from aggregated case data, we applied a disaggregation regression model trained on subnational case reports from 14 countries in Central and South America, using data from 2018 and 2019 to maximise spatial coverage and resolution. The model uses environmental and demographic covariates to downscale aggregated case counts to ∼10 km^2^ resolution. Model performance was evaluated using 5-fold spatial block cross-validation, wherein the study area was divided into five spatial blocks of approximately equal size. We selected five spatial blocks to balance spatial representativeness with computational feasibility, as increasing the number of folds would have substantially raised the computational burden given the scale and resolution of the dataset. The model was iteratively trained on four blocks and tested on the fifth to assess predictive accuracy within the training region. The reported correlation reflects the agreement between observed and model-predicted dengue incidence at the administrative level, calculated by aggregating pixel-level model predictions and comparing them to reported surveillance data. We used Spearman’s correlation because it assesses monotonic relationships and is more robust to non-normal distributions and outliers, making it suitable for our data, which includes skewed incidence values and varying case magnitudes across regions [27,28]. The posterior means and 95% credible intervals for the estimated covariate effects and hyperparameters are presented in supplementary **Figure S1**. Temperature and urban built-up areas were positively associated with dengue incidence, while normalized difference vegetation index (NDVI) and humidity were negatively associated. Spatial hyperparameters confirmed the presence of strong residual spatial structure, supporting the inclusion of spatial random effects in the model.

The model achieved a mean incidence correlation (Cor_I) of 0.72, calculated as the Spearman correlation between observed and predicted incidence in the selected countries of South America, suggesting moderately strong predictions within the training region. **Figure 1 A&B** illustrates the transition from aggregated dengue case data to the resulting fine-scale predicted cases through the disaggregation regression framework. Figure 1A shows the reported dengue incidence per 100 000 inhabitants, collected at administrative levels 1 and 2 across Central and South America, highlighting areas with high reported burden, particularly in Brazil, Colombia, Mexico, and Venezuela. These data form the basis for model fitting, yet their spatial resolution is limited by the reporting standards in each country. Figure 1B shows the predicted dengue cases, modeled at ∼10 km^2^ resolution. High case concentrations are predicted in densely populated areas such as southeast Brazil, northern South America, and parts of Central America, while lower predicted case counts are observed in the southern cone.

**Figure 1:**
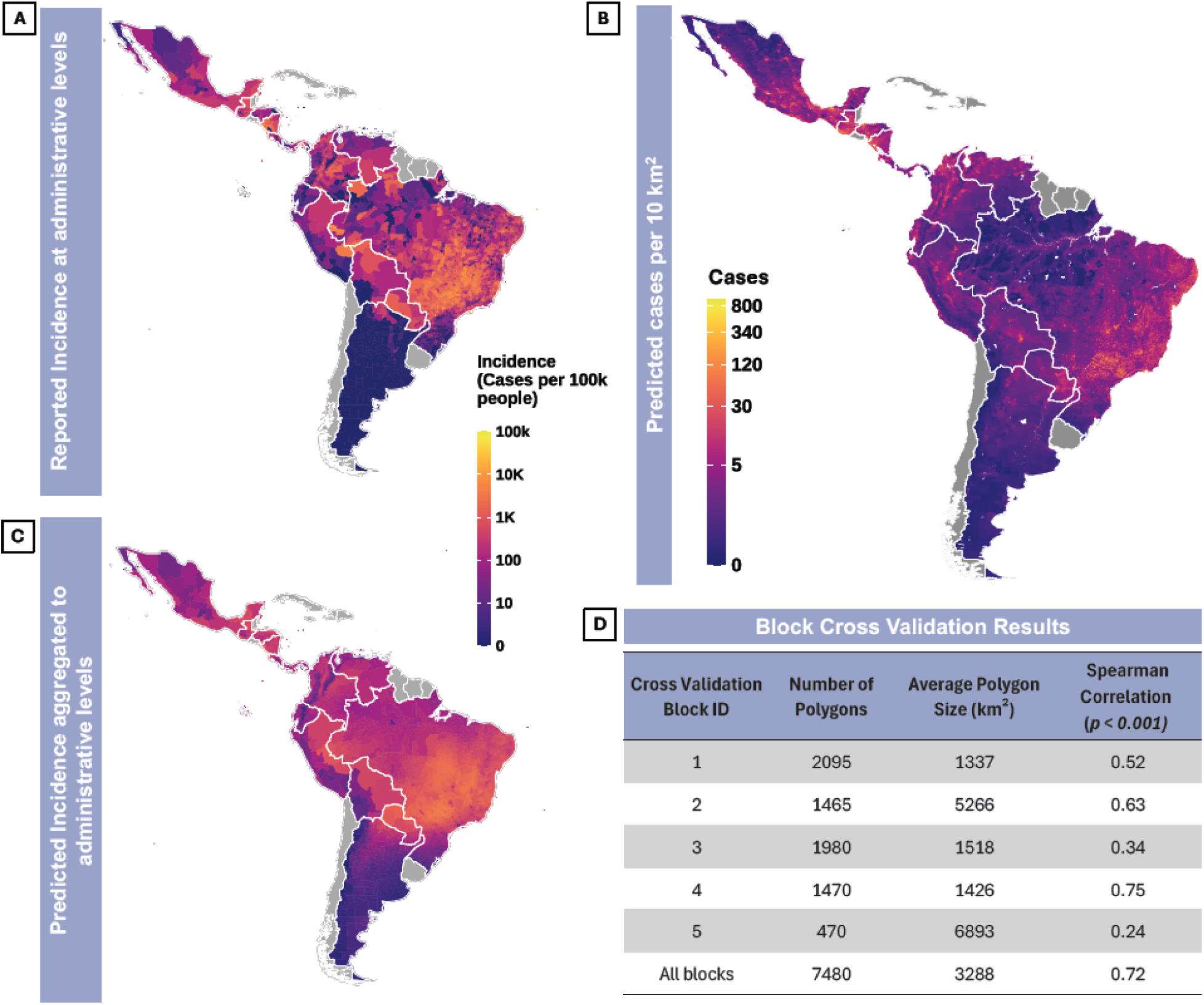
**From reported dengue cases to model-predicted incidence across Central and South America**. Panel (A) shows aggregated reported dengue case counts at administrative levels 1 and 2 for the year 2019, used as input for model fitting. Panel (B) displays predicted cases derived from the disaggregation regression model at ∼10 *km*^2^ resolution, capturing fine-scale spatial variation in dengue burden for that year. Panel (C) presents the predicted incidence (per 100k inhabitants), calculated by aggregating the predicted cases to administrative level 1 and 2 and normalising by population size. Panel (D) presents the results from the 5-fold block cross validation.

To evaluate model generalizability, the model was then trained on all five blocks and predicted onto seven countries (Australia, India, Malaysia, Myanmar, Phillipines, Saudi Arabia and Thailand) for which dengue incidence data were available at least at the first administrative level outside of Central and South America. Supplementary **Figure S2** shows scatter plots comparing predicted and observed dengue incidence at the administrative level, with correlation used as evaluation metric, as it reflects per capita risk rather than absolute burden. The strongest positive correlation was observed in Myanmar (Cor_I = 0.76, p = 0.0061), followed by Australia (Cor_I = 0.51, p = 0.0062) and Thailand (Cor_I = 0.25, p = 0.0031), suggesting reasonable alignment between predictions and reported data in these settings. In contrast, Malaysia exhibited a significant negative correlation (Cor_I =-0.22, p = 0.0519), while correlations were weak in India (Cor_I = 0.26, p = 0.0121) and Philippines (Cor_I = 0.11, p = 0.0394) or non-significant in Saudi Arabia (Cor_I= 0.06, p = 0.0914). Those poor correlations may reflect a combination of factors, including differing population immunity profiles, localized vector control interventions, or inconsistencies in surveillance and reporting systems that affect the comparability of observed data across regions.

To further interpret model performance and contextualize the observed correlations, we compared reported incidence to three additional dengue risk indicators: transmission potential, vector (*Aedes aegypti*) suitability, and dengue suitability (**Supp Figure S3**). For each of the seven independent countries, we extracted these indicators by averaging the values of each risk surface over the corresponding administrative units. Transmission potential also exhibited relatively strong alignment in several countries, including Myanmar (Cor = 0.67, p = 0.001) and Australia (Cor = 0.51, p = 0.01), but was weak and insignificant in others such as Malaysia (Cor =-0.06, p = 0.08). *Aedes* suitability showed moderate correlations in Myanmar (Cor = 0.49, p = 0.005) and India (Cor = 0.5, p < 0.001). Dengue suitability, while positively correlated in Myanmar (Cor = 0.71, p < 0.001) and India (Cor = 0.57, p < 0.001), displayed weak or negative associations in countries like Thailand (Cor = –0.15, p = 0.02) and Philippines (Cor = –0.11, p = 0.03). These findings highlight the challenges of transferring models across regions with differing epidemiological dynamics, surveillance systems, and population distributions.

### Predicting Dengue Incidence in Africa

The trained disaggregation regression model was then applied to the African continent to generate high-resolution predictions of dengue incidence based on environmental and demographic covariates for Africa. **Figure 2A** shows predicted cases per 10km^2^ in Africa, the map highlights regions of west Africa as high-burden zones, a pattern driven primarily by high incidence rate combined with large population sizes and recent urban growth. The predicted pixel-level incidence map (**Supp Figure S4A**) reveals a heterogeneous distribution of dengue risk across Africa, with areas of elevated incidence concentrated across the Sahel region of Africa (Senegal, Mauritania, Mali, Burkina Faso, Niger, northern Nigeria, Chad, Sudan, and Eritrea) and parts of eastern Ethiopia and Somalia. These spatial patterns appear to align with areas where conditions may be increasingly favorable for dengue transmission [29]. In contrast, lower predicted incidence are observed in northern and southern Africa, regions characterized by climatic conditions generally considered less suitable for *Aedes* mosquito survival and dengue virus transmission [30]. We can further observe this pattern looking at the aggregated dengue incidence per country (**Figure 2B**). District-level (administrative level 1) aggregation highlights subnational variation, revealing localized hotspots; particularly in Sudan, Ethiopia, and Senegal (**Figure 2C** and **Supp Figure S5**). We also show model uncertainty (**Supp Figure S4B**) associated with the predicted incidence across Africa and we find that much of the variation is concentrated in the Sahel region.

**Figure 2:**
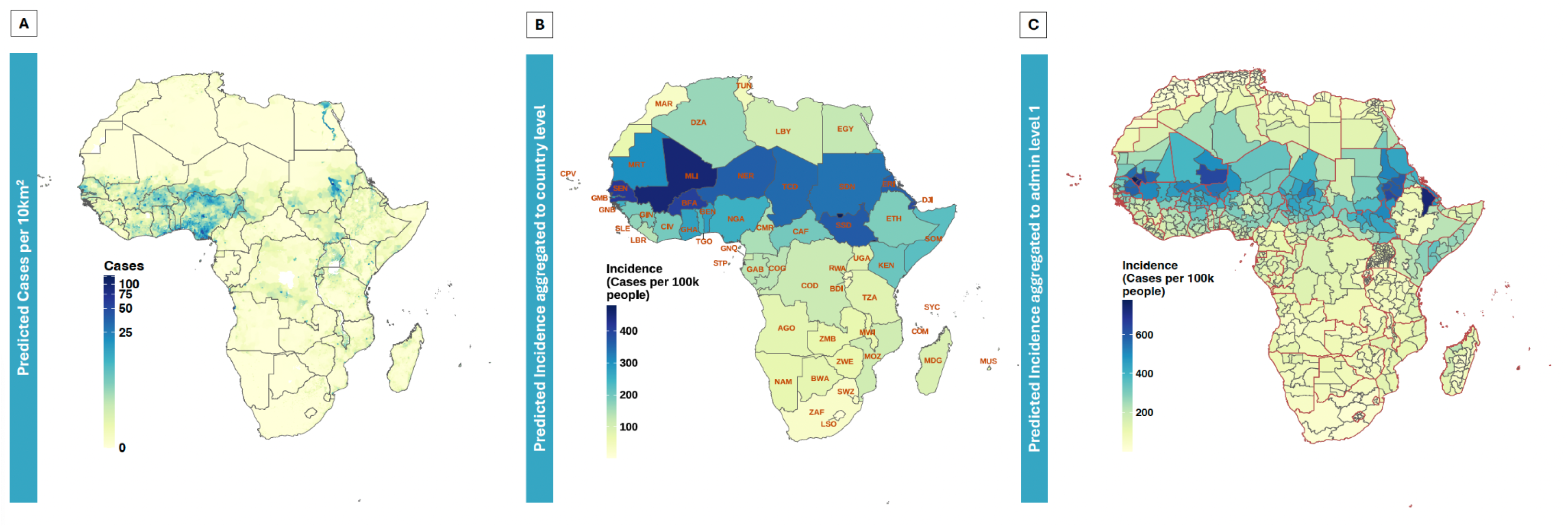
**Spatial predictions of dengue cases and incidence across the African continent using the disaggregation regression model trained on case data from Central and South America**. Panel (A) displays predicted cases at pixel-level (10km^2^). Panel (B) shows the incidence at country level. Panel (C) shows the incidence at administrative level 1 in Africa. AGO = Angola, BEN = Benin, BFA = Burkina Faso, BWA = Botswana, CAF = Central African Republic, CIV = Côte d’Ivoire, CMR = Cameroon, COD = Democratic Republic of the Congo, COG = Republic of the Congo, COM = Comoros, CPV = Cabo Verde, DJI = Djibouti, DZA = Algeria, EGY = Egypt, ERI = Eritrea, ETH = Ethiopia, GAB = Gabon, GHA = Ghana, GIN = Guinea, GMB = Gambia, GNB = Guinea-Bissau, GNQ = Equatorial Guinea, KEN = Kenya, LBR = Liberia, LBY = Libya, LSO = Lesotho, MDG = Madagascar, MLI = Mali, MOZ = Mozambique, MRT = Mauritania, MUS = Mauritius, MWI = Malawi, NAM = Namibia, NER = Niger, NGA = Nigeria, RWA = Rwanda, SDN = Sudan, SEN = Senegal, SLE = Sierra Leone, SOM = Somalia, SSD = South Sudan, STP = São Tomé and Príncipe, SWZ = Eswatini, SYC = Seychelles, TCD = Chad, TGO = Togo, TUN = Tunisia, TZA = Tanzania, UGA = Uganda, ZAF = South Africa, ZMB = Zambia, ZWE = Zimbabwe.

To explore where model predictions and reported observations align or diverge, we compared them to country-level reported (suspected) dengue cases compiled in a review by [31], to which we added cases reported for the year 2024 compiled from publicly available reports. The resulting correlation (**Figure 3A**) was moderate (Cor_I = 0.33, p = 0.01), indicating a reasonable alignment in the ranking of countries by predicted and reported dengue burden. Several countries; including Ivory Coast (CIV), Comoros (COM) and Togo (TGO) showed close alignment between predicted and reported incidence. Others such as Mauritius (MUS) and Sao Tome and Principe (STP), the model underpredicted incidence in these countries relative to reported case numbers. Nigeria exhibits high predicted dengue incidence alongside elevated dengue suitability, vector suitability, and transmission potential. However, reported case numbers remain comparatively low. This divergence could likely reflect underreporting or gaps in surveillance rather than true absence of disease, which highlights areas that require enhanced diagnostic capacity and more robust surveillance infrastructure.

**Figure 3:**
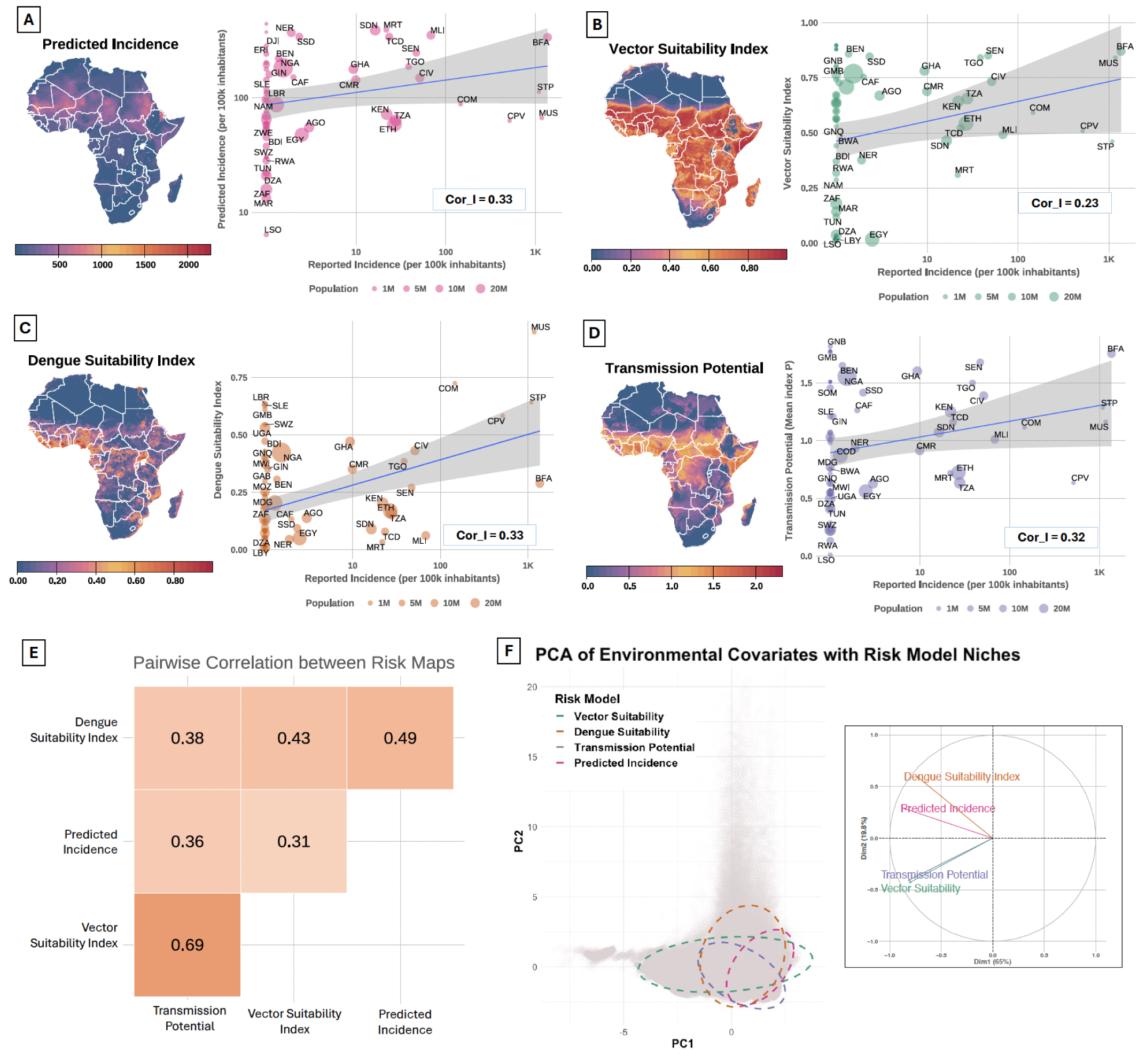
Comparison of four dengue risk indicators across Africa. Panel (A-D) Comparison of transmission potential, vector suitability, dengue suitability, and predicted incidence, against reported suspected dengue incidence at the country level. Correlation values (Cor_I) reflect the strength of association between each model and reported incidence. Circle size shows country population. Panel (E) Pairwise correlations between four dengue risk maps across Africa. Panel (F) Principal Component Analysis (PCA) of the environmental covariate space (temperature, humidity, NDVI, GDP, and urbanization) across Africa, overlaid with ellipses representing the spatial footprint of each dengue risk indicator. The distribution of points reflects the range of environmental conditions across the continent, while the ellipses indicate the environmental domains emphasized by each model. Panel (F-Inset) PCA biplot showing the spatial relationships among four dengue risk models. Arrows represent the direction and strength of each model’s contribution to the overall variation in risk patterns. Models pointing in similar directions exhibit more similar spatial distributions.

We also assessed the relationship between the three additional dengue risk indicators and country-level suspected dengue cases in Africa (**Figure 3)**. Along with our predicted incidence model, dengue suitability index (**Figure 3C**) and Transmission potential (**Figure 3D**) showed the strongest positive correlations with reported cases (Cor_I = 0.33 and 0.32, respectively) and vector suitability (**Figure 3B**) showed the weakest association (Cor_I = 0.22). In Burkina Faso and Senegal, while the virus-based environmental suitability model predicts relatively low suitability on average, both the vector suitability and transmission potential indices indicate elevated values. This suggests that although ecological conditions specifically favoring dengue transmission may be limited according to the dengue suitability model, the broader suitability for the vector and its conducive transmission conditions are present, highlighting a potential underestimation of dengue risk when relying solely on virus-based environmental suitability.

To examine the degree of spatial agreement between all four dengue risk indicators, we computed pixel-wise correlations (cor) between all pairs of risk maps (**Figure 3E**). The highest correlation was observed between the Transmission Potential and Vector Suitability Index (cor = 0.69, p<0.001), likely due to their shared reliance on temperature and humidity. The correlations between the predicted incidence and other models are moderate; cor = 0.36 (p<0.001) with transmission potential, cor = 0.31 (p<0.001) with vector suitability, and cor = 0.49 (p<0.001) with the dengue suitability index. The dengue suitability index, derived from presence-only data, showed the weakest overall correlations with other surfaces, potentially due to differences in occurrence data quality, spatial resolution, or bias in reporting locations. The Principal Component Analysis (PCA) of the environmental covariates used in this study (temperature, humidity, NDVI, GDP and urbanisation) revealed the distinct environmental niche occupied by each of the four dengue risk models (**Figure 3F).** In the main plot, ellipses indicate the environmental space where each model’s high predictions are concentrated. Transmission potential and vector suitability models occupy similar regions of environmental space, indicating strong alignment with climatic variables such as temperature and humidity. In contrast, the predicted incidence and dengue suitability models show more distinct and less overlapping distributions, potentially suggesting a stronger association with socio-environmental variables like GDP and urbanisation. The inset panel (**Figure 3F**) presents a PCA of the four risk model outputs, summarizing their spatial correlation structure across the study area. The PCA biplot shows that transmission potential and vector suitability align with a similar underlying gradient, while predicted incidence and dengue suitability are oriented along a different gradient.

### Overlap Between Model-Predicted Incidence and Other Dengue Risk Indicators in Africa

To assess how our model qualitatively compares with existing risk frameworks, we compared the spatial estimate output from our disaggregation model: predicted incidence (**Figure 3A**) and three widely used dengue risk indicators: (i) the dengue Suitability Index (**Figure 3C**), (ii) the transmission potential (Index *P*) (**Figure 3D**), and (iii) the vector suitability index for *Aedes aegypti* (**Figure 3B**). Assessing agreement and divergence among them provides insight into where models converge to flag consistently high-risk areas, and where uncertainties remain.

To enable direct spatial comparison and identify areas of high-risk concordance, we applied model-specific thresholds to each risk surface: >0.5 for both the *Aedes* vector suitability and dengue environmental suitability indices, >0.5 for the transmission potential (Index P) [9], and >1 predicted case (per 100k inhabitants per pixel) for the disaggregation-derived incidence map (**Figure 4-Inset**). We then created binary high-risk maps and overlaid them to count how many models flagged each pixel as high-risk (**Figure 4**). The comparison between all four dengue risk maps reveal high concordance across much of the coastline of West Africa, in countries such as Nigeria, Benin, Togo, Ghana and Ivory Coast. We also observe notable overlap across parts of the southern Sahel (including Burkina Faso, southern Mali, and southwestern Niger, northern part of Nigeria). In these regions, the predicted incidence from the disaggregation model aligns spatially (**Figure 4-Inset**) with both high transmission potential and vector suitability, suggesting that environmental and epidemiological conditions are simultaneously favorable for dengue transmission. Mauritius also seems to be at the intersection of high risk across at least three indicators.

**Figure 4:**
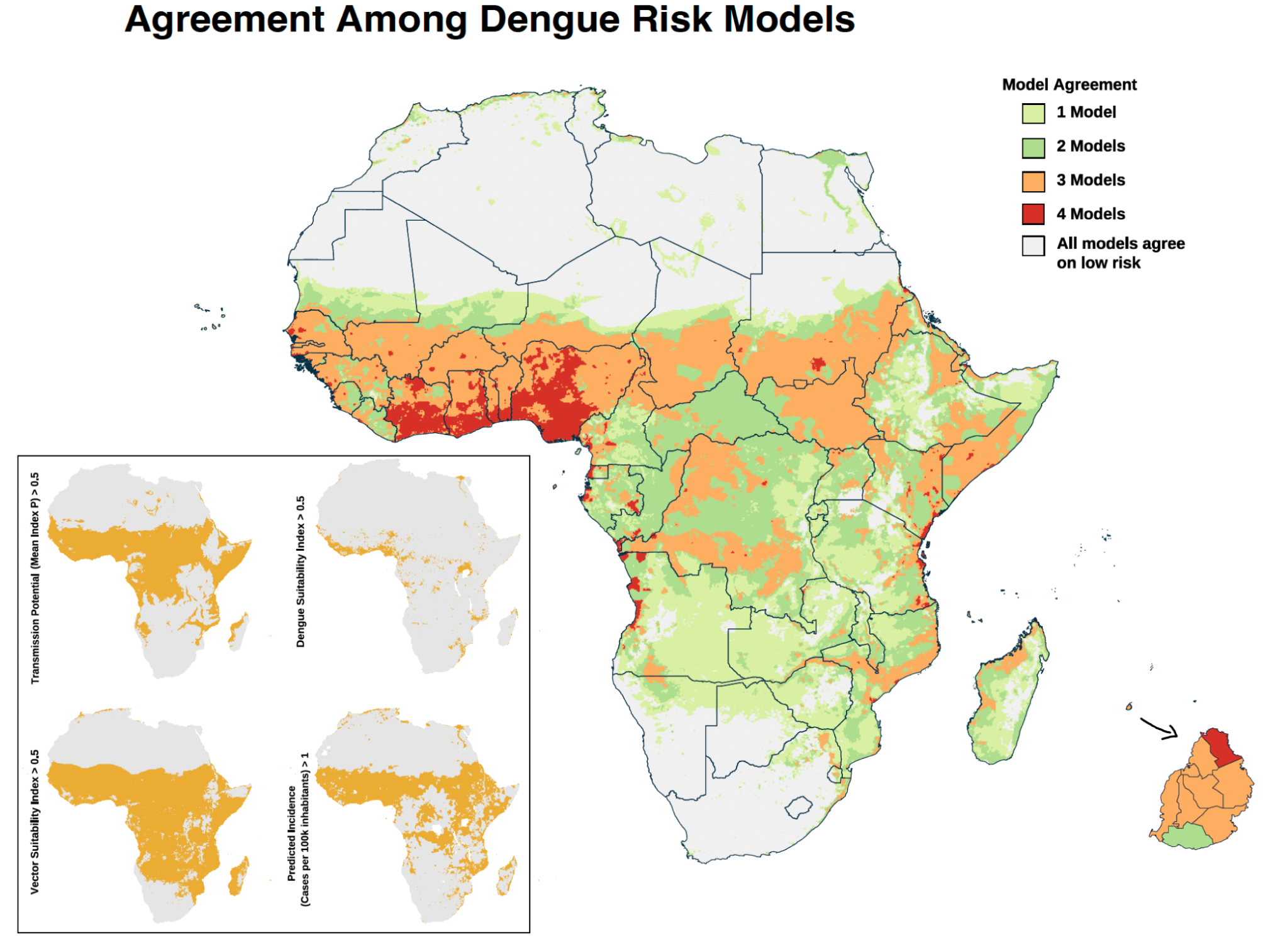
Agreement among dengue risk models across Africa. Map showing the number of dengue risk indicators: predicted incidence, transmission potential, dengue suitability index and vector suitability index, that classify each pixel as high risk. High-risk classification was determined using model-specific thresholds: >0.5 for suitability indices and transmission potential; >1 of predicted incidence per 100k people (per pixel). Darker red areas indicate strong agreement among all models in predicting high dengue suitability, whereas lighter green areas reflect lower concordance. Grey regions represent areas where all four models predicted low risk based on the respective threshold.

In contrast, several areas of Eastern Africa and Central Africa exhibit elevated transmission potential and vector suitability but relatively limited overlap with the other risk maps. These mismatches may reflect underlying differences in model assumptions, such as the emphasis on ecological suitability versus actual disease burden, or differences in input datasets, including climatic baselines and mosquito surveillance data. From a public health standpoint, such discrepancies between model types underscore the importance of not overlooking regions simply because ecological models signal lower risk. Instead, these areas may represent critical frontiers for proactive surveillance and risk validation, particularly in the face of rapid urban growth and climate variability.

## Discussion

This study presents the first continent-wide estimates of subnational dengue burden in Africa using a data-driven, incidence-based modeling framework. By applying disaggregation regression to coarsely aggregated case data, we demonstrate its utility in generating fine-scale, interpretable predictions of dengue incidence; information that is directly tied to disease burden and highly relevant for public health planning. In contrast to vector or occurrence-based models, which reflect ecological suitability or presence, disaggregation regression emphasizes transmission intensity and expected case load. Crucially, our comparison of the three widely used risk mapping approaches alongside the disaggregation-derived predicted incidence reveals that these models capture different dimensions of risk and often diverge in the areas they flag as high priority. Rather than viewing these outputs as competing or conflicting, our findings underscore the importance of using them in a complementary manner to inform more robust and context-sensitive decision-making, especially in data-scarce settings like Africa.

Within the training region, the disaggregation model achieved moderate predictive accuracy (mean correlation = 0.72), capturing spatial variation in dengue burden at subnational levels. Importantly, the model preserved spatial heterogeneity in risk when downscaling to ∼10 km^2^ resolution, allowing the identification of localized predicted high-incidence clusters that would otherwise be obscured in coarser aggregated data. Across the seven countries assessed, we observed strong correlation at administrative level 1 in Myanmar and Australia (**Figure S2**). We also found varying degrees of alignment between reported dengue cases and each of the four risk indicators at administrative level 1. Predicted incidence and transmission potential both displayed a negative correlation with reported cases in Malaysia while vector and dengue suitability displayed a positive correlation (**Figure S3).** These results point towards the importance of considering local epidemiological contexts, health system capacities, and surveillance completeness when applying incidence-based or other risk models across regions. In particular, poor performance in some countries may reflect substantial underreporting, limited diagnostic capacity, or inconsistencies in case definitions; all of which reduce the reliability of input data and constrain the model’s ability to capture true transmission patterns [32,33].

Despite known challenges in building disaggregation regression models [34]; particularly in data-scarce settings or when working with coarsely aggregated case data, this approach remains a valuable addition to the dengue risk mapping toolkit. While performance may be sensitive to the scale and quality of input data, especially under model misspecification [25,35], these models offer a unique opportunity to translate available incidence data into interpretable, high-resolution surfaces. Rather than replacing other frameworks, disaggregation regression helps uncover potential transmission dynamics that may be overlooked by purely ecological or mechanistic approaches, reinforcing the value of combining multiple perspectives in areas where data limitations are the norm.

Moreover it is also important to examine both metrics: predicted incidence (**Figure 3A**) and total predicted cases (**Figure 2A**) to gain a comprehensive understanding of dengue risk. Incidence reflects the intensity of transmission, identifying areas where individuals face the highest likelihood of infection. This is particularly useful for detecting emerging hotspots and targeting interventions aimed at reducing transmission risk [36,37]. In contrast, predicted cases are driven by both incidence and population size, offering insight into the potential burden on health systems [38]. Relying on only one of these metrics may lead to an incomplete or misleading assessment of risk; together, they provide a more nuanced picture that supports both targeted transmission control and broader resource allocation planning.

While risk projections inherently comprise large margins of uncertainty; given the lack of training data from Africa, they provide a starting point for further investigation and surveillance planning. The model identified several areas with elevated risk, including coastal West Africa and the Congo Basin, and parts of Eastern Africa (**Figure 2A**). These predictions align broadly with regions previously suggested to be vulnerable based on vector presence and climatic suitability [39,40]. Subnational estimates **(Figure 2C)** revealed localised pockets of elevated incidence, for instance, the 2023 outbreak in Ethiopia started in the Afar region (**Figure S5**) [41]. The predicted incidence for Burkina Faso also followed closely patterns found in [42] (**Figure S5**).

Moreover, our results were in line with case numbers reported in the review by Mwanyika et al., (2024) and the perspective piece by [20]. The moderate correlation between model-predicted case counts and reported suspected dengue cases (2013-2024) (Cor_I = 0.32) suggests that the model reasonably captures the relative distribution of dengue burden across countries, despite the lack of African data in model training. The correlation between the other three dengue risk indicators were also modest, with the highest correlation of 0.33. The weak correlations may reflect differences in reporting systems, or surveillance coverage, which can influence case detection independently of true transmission intensity.

Several methods have been developed to improve understanding of the spatial distribution of dengue burden and to identify areas at elevated risk, with the aim of informing burden estimation, enhancing preparedness, and guiding public health response strategies. Each method carries its own set of assumptions and limitations, offering a distinct perspective on dengue risk. Whether focused on ecological suitability, transmission potential, or estimated disease burden; these approaches emphasize the importance of understanding what type of information each approach is designed to capture [4]. Comparisons between the disaggregation model and three widely used dengue risk indicators (vector suitability index, dengue suitability index, and transmission potential) revealed areas of both concordance and divergence. Overlapping high-risk zones in coastal West Africa suggest a strong signal of ecological and epidemiological suitability across models (**Figure 4**). The central region of Burkina Faso shows overlap across all four models, indicating a potential high-risk area that aligns with regions previously affected by major dengue outbreaks [42]. We also observe overlaps across all four dengue risk indicators in regions of Eastern Africa, notably the coastal part of Kenya and Tanzania where dengue circulation has previously been reported [43].

However, across most parts of central Africa we observe fewer model agreements which highlight the differing assumptions, inputs, and outputs of these models (**Figure 3F**) [29,44]. We observe that vector suitability and transmission potential occupy more similar regions of environmental space compared to the dengue suitability and predicted incidence models. For example, the vector suitability index assumes that areas with suitable temperature and humidity for *Aedes aegypti* are at high risk, which leads it to flag much of Angola and Zimbabwe as high-risk, where dengue occurrence data are lacking. In contrast, the disaggregation model captures actual disease burden as reflected in reported cases and is thus more directly epidemiological in nature. Moreover we identified regions where only our incidence-based model predicted elevated dengue risk (**Supp Figure S6**), including countries such as Burundi and Rwanda. In Rwanda, for example, our model estimates approximately 735 cases annually. Notably, a recent seroprevalence study reported that 30.4% of 2,286 samples tested positive for dengue antibodies[45], with the highest proportion of seropositive individuals observed in the eastern region, an area that our model also highlights as having the greatest predicted burden (**Supp Figure S5**).

These divergences between approaches are not necessarily flaws but reflect the fact that different approaches intentionally seek to model distinct facets of transmission risk. In settings, where surveillance is limited or ecological data are sparse, model selection is often driven by data availability and end-user needs. While ecological niche models may highlight potential for vector presence, mechanistic models describe climate-based thresholds for transmission, and disaggregation regression translates case data into spatial burden estimates. Depending on the question at hand; whether estimating current burden, forecasting future trends, or targeting intervention, different tools may be more appropriate[14]. Importantly, the answers to complex global health questions may emerge from triangulating across multiple modeling frameworks, even when they yield contrasting results.

Despite its contributions, this study has several limitations. First, the disaggregation model was trained exclusively on Latin American data and applied to Africa, which may limit its accuracy in new ecological and epidemiological contexts. There are modelling avenues that need to be further explored, especially with dengue being most common in urban settings. Another key limitation is that we estimate annual incidence, which may obscure important temporal variation and mask periods of heightened transmission risk, given the seasonal nature of dengue outbreaks. The quality and granularity (e.g size of polygons/districts) of input case data varied by country, and underreporting which is common in dengue surveillance may also introduce bias. The comparisons with other risk models also rely on static surfaces and may not reflect dynamic changes in climate or transmission potential over time.

In conclusion, disaggregation regression offers a promising tool for generating high-resolution, incidence-based dengue risk maps in data-limited settings. Despite the challenges of cross-regional application, the model successfully captured meaningful spatial patterns in dengue burden and produced predictions for Africa that are broadly consistent with the distribution of reported cases to date. Our comparative analysis underscores the importance of considering both ecological suitability and epidemiological burden when mapping dengue risk, particularly in under-studied regions such as Africa. Future work should be built by incorporating multi-model approaches, improving real-time data availability, and expanding ground-truth case reporting to further strengthen public health preparedness and response.

## Materials and Methods

### Dengue Case Data

We compiled dengue case data from 14 countries in Central and South America where subnational surveillance records are available for the year 2019, while using data from a single year does not allow for assessment of temporal generalizability, it enables a focused evaluation of the model’s spatial prediction capacity. These countries are: Argentina, Bolivia, Brazil, Colombia, Costa Rica, Ecuador, Guatemala, Honduras, Mexico, Nicaragua, Panama, Paraguay, Peru, and Venezuela. We obtained case data from a combination of sources, including national ministries of health, the Pan American Health Organization (PAHO), and other publicly available government epidemiological bulletins and archives. The spatial resolution of the data varies by country: most case records were reported at the first administrative level (admin1) (e.g., provinces or departments), while finer-resolution data at the second administrative level (admin2) (e.g. municipalities) were available for selected countries such as Brazil and Argentina.

To test the generalizability of the model to other regions, we obtained independent dengue case data for seven countries (testing regions) located outside the initial training region, namely: Australia, India, Malaysia, Myanmar, Philippines, Saudi Arabia and Thailand. We selected these countries based on the availability of administrative-level 1 data and because they represent diverse ecological and epidemiological settings. We used this independent data exclusively for out-of-sample evaluation.

### Environmental and Socioeconomic Covariates

We assembled a suite of environmental and socioeconomic covariates hypothesized to influence dengue transmission, based on extensive literature linking these variables to vector ecology, viral dynamics, and host exposure. The covariates include: temperature, precipitation, humidity, Normalized Difference Vegetation Index (NDVI), urbanisation, Gross Domestic Product (GDP) (as a proxy for socioeconomic development and access to health services) [46–49] (**Supp Figure S7**). We harmonized all covariates to a common spatial resolution (∼10 km²) and time period (2019) to align with the case data. Climatic variables were obtained from Climate Data Store [50] https://cds.climate.copernicus.eu/datasets/ecv-for-climate-change?tab=overview, NDVI from NASA (https://modis.gsfc.nasa.gov/data/dataprod/mod13.php), and urbanization and GDP layers from EarthEnv (https://www.earthenv.org/landcover) and https://zenodo.org/records/13943886 [51]. Where necessary, we aggregated monthly raster layers to an annual mean.

### Disaggregation Regression Modelling framework

To estimate fine-scale dengue incidence from polygon-level case reports, we implemented a disaggregation regression model using the disaggregation R package [52], which is based on the framework developed by [53] and related spatial modeling literature [25,35]. This approach downscales aggregated case counts into a continuous pixel-level incidence rate surface by leveraging spatially referenced covariates and incorporating spatial dependence.

Let *Y_A_* denote the observed case count in polygon A, with total population *N_A_*, composed of a set of pixels *p ɛ A*. We assume that

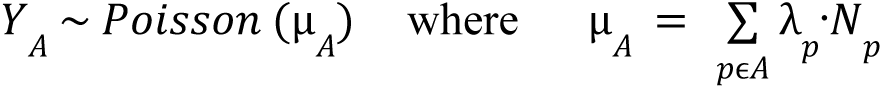

where λ*_p_* is the latent incidence rate at pixel *p* and *N_p_* is the population count associated with pixel *p*.

The log incidence rate at each pixel is modeled as:

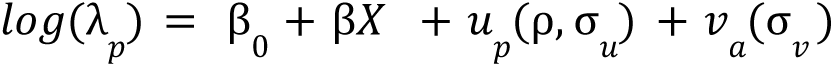

where:

- β_0_ is the intercept
- *X_p_* is the vector of covariates
- β is the vector of slope coefficients
- *u_p_* is a spatially structured random effect modelled as a Gaussian random field
- *v_a_* is a polygon-level independent and identically distributed (iid) random effect to account for unstructured spatial variation and models overdispersion in the data.

The spatial field *u_p_* follows a Matérn covariance function, defined by two hyperparameters: ρ, the nominal spatial range and σ*_u_* the marginal standard deviation that controls the overall variability of the spatial field. The iid random effect captures both unobserved explanatory variation and excess variability not accounted for by the Poisson model. We constructed the spatial mesh using finite elements, adhering to standard mesh-building guidelines from the INLA framework [54] to balance computational efficiency and spatial resolution.

The disaggregation model relies on Template Model Builder (TMB) within the disaggregation R package, which uses the Laplace approximation to estimate model parameters. We selected prior distributions for the model parameters based on practical insights from previous applications of disaggregation regression, particularly the framework outlined in Lucas et al. (2022)[25]. We complete the model by setting priors n the parameters β_0_, β, ρ, σ_u_, σ*_v_*. The spatial field parameters ρ and σ, we applied joint penalised complexity priors that favour smooth spatial fields. We also assigned σ*_v_* a penalised complexity prior such that P(σ*_v_* >0.2) = 0.05. This choice was based on a comparison between the variance expected under a Poisson model and plausible bounds for case uncertainty due to under-reporting and case clustering, as described in Lucas et al. (2021) [25]. Finally we set the priors on the regression coefficient, β ∼ Norm (0,1) allowing for flexibility in covariate effects.

The final output includes two risk maps:

1. Incidence rate map (λ*_p_*) predicted per capita dengue risk at ∼10 km² resolution.
2. Incidence map: computed by multiplying incidence rate by pixel-level population (λ*_p_.N_p_*) yielding predicted case counts per pixel per year.

### Model Evaluation

We evaluated the performance of the incidence-based model by calculating the polygon-level Spearman correlation coefficient between the observed and predicted aggregated incidence rates using two approaches: (i) *k*-fold spatial block cross-validation within the training region (Central and South America), where the region was divided into five spatial blocks of approximately equal size, and the model was iteratively trained on four blocks and tested on the remaining one and (ii) evaluation using the unseen data from countries outside the training data set; the model was trained on the entire set of five spatial blocks and then tested on these external countries. In both cases, the model’s high-resolution predictions were aggregated to the administrative unit level for direct comparison with surveillance-reported case data.

The model produces a continuous spatial landscape of predicted incidence rates (per pixel). To estimate the total case burden, we multiplied the predicted incidence rate by the population count at each pixel level; representing predicted case counts per pixel. For each administrative polygon or country, we calculated dengue incidence per 100,000 inhabitants by first summing the predicted pixel-level case counts within each polygon and then dividing by the total population of that polygon. To assess model performance, we then calculated correlation metrics: (1) Cor_I: the correlation between the total reported incidence and the predicted incidence at administrative level 1 and country level in the case of Africa where only national level data were available.

### Continental Prediction and Reaggregation for Africa

In addition to generating predictions for the countries within the training and testing regions, we applied the disaggregation model to the African continent using the same covariate layers prepared for model training. Our model produces pixel-level incidence rate estimates at 10 km² resolution across Africa. To facilitate comparison with available national-level dengue data, we reaggregated these pixel-level predictions to the country level by summing the predicted number of cases across all grid cells within each country. This reaggregation allowed for an indirect comparison between predicted national dengue burdens and officially reported country-level case counts, which provides an additional layer of external face validation in the absence of widespread subnational surveillance data across Africa.

### Comparing Dengue Risk Maps for Africa

To contextualize our model’s outputs, we compared our predicted incidence map to three established dengue risk mapping approaches:

1. **Transmission Potential** Map (Index P): We obtained spatio-temporal estimates of climate-based dengue transmission potential, referred to as Index P, from Nakase et al. (2023)[26]; https://doi.org/10.6084/m9.figshare.21502614). This index quantifies the transmission capacity of a single adult female Aedes mosquito over its lifetime within a fully susceptible human population. It integrates temperature-dependent biological parameters, including the mosquito’s lifespan, biting rate, oviposition frequency and incubation periods.
2. **Dengue Suitability Index** (Occurrence-based ENM): We used the dengue suitability index produced by Lim et al. (2025) [6]. The authors applied an ecological niche modeling approach trained on global dengue occurrence data and a range of environmental, climatic, and socioeconomic covariates known to influence transmission were used as model inputs. This index reflects the environmental suitability for the virus itself, based on historical patterns of reported dengue presence.
3. *Aedes aegypti-* **Vector Suitability Index**: A global ecological niche model of *Aedes aegypti* distribution which indicates the environmental suitability for the primary dengue vector [5]. While this is not a direct measure of dengue suitability, such vector-based maps are often used as proxies for dengue risk, given the central role of *Aedes aegypti* in transmission. Throughout the manuscript, we include it alongside the other three risk layers transmission potential, dengue suitability, and predicted incidence and refer to all four collectively as dengue risk indicators.

We scaled and aligned all risk maps to a common spatial resolution (∼10 km²) and extent (Africa) for direct spatial comparison. To examine the degree of spatial agreement between all four dengue risk indicators, we computed pixel-wise correlations (cor) between all pairs of risk maps and to visually assess the agreement between maps, we used a combination of visual inspection, bivariate overlay maps, analysis of covariate response patterns and principal component analysis (PCA). Furthermore, to systematically evaluate spatial concordance across the four frameworks, we selected threshold values to each risk surface: >0.5 for both the *Aedes* vector suitability and dengue suitability indices, >0.5 for the transmission potential (Index P) [9], and >1 predicted case for the disaggregation-derived incidence map.

These thresholds were selected to identify pixels classified as high-risk according to each model’s individual scale and interpretation of dengue risk.

Binary (high-risk/not high-risk) surfaces were then generated for each map based on these thresholds, and overlaid to determine the number of models in agreement for identifying a given pixel as high-risk. Pixels were categorized from 1 to 4 depending on how many models exceeded their respective thresholds at that location. This approach enabled the detection of areas with consistent agreement across models, as well as those flagged by only a subset of risk mapping frameworks. By focusing on the spatial convergence of independently derived risk surfaces, this method highlights regions of overlapping high risk and offers insights into the robustness and complementarity of different modeling approaches.

## Data Availability Statement

All dengue case data used in this study, as well as the environmental covariates and model outputs, are available on the project’s GitHub repository:

https://github.com/CERI-KRISP/Dengue_Incidence_Prediction.git. This includes the aggregated case data used for model training, the raster layers used as covariates, and the resulting predicted incidence maps. In addition, the alternative dengue risk maps used for comparison (vector suitability, dengue suitability, and transmission potential) are also provided in the repository for full reproducibility.

## Contributors

J.P., H.T and M.D conceptualised and designed the study. J.P. analysed data, executed all primary data visualizations, and wrote the original draft. J.P and G.M. collected and curated epidemiological data. J.P, T.L. and H.T interpreted data. T.L, J.L and M.U.G.K. contributed to data analysis and visualisation. T.d.O and H.T. acquired funding for this project. H.T., M.D., T.d.O supervised the study. All authors had full access to all the data in this study. All authors reviewed and edited the final draft. All authors had final responsibility for the decision to submit for publication.

## Acknowledgements

The authors acknowledge that this work is an initiative of the CLIMADE consortium. The Centre for Epidemic Response and Innovation and the Kwazulu-Natal Research Innovation and Sequencing Platform are supported in part by grants from the Rockefeller Foundation (HTH 017), and the INFORM Africa project through the Institute of Human Virology Nigeria (U54 TW012041), Global Health EDCTP3 Joint Undertaking and its members and the Bill & Melinda Gates Foundation (101103171), the Health Emergency Preparedness and Response Umbrella Program, managed by the World Bank Group (TF0B8412), the Medical Research Foundation (MRF-RG-ICCH-2022-100069), the Wellcome Trust (228186/Z/23/Z), and the Novo Nordisk Foundation (NNF24OC0094346). M.U.G.K. acknowledges funding from The Rockefeller Foundation (PC-2022-POP-005), Google.org, the Oxford Martin School Programmes in Pandemic Genomics & Digital Pandemic Preparedness, European Union’s Horizon Europe programme projects MOOD (#874850) and E4Warning (#101086640), Wellcome Trust grants 303666/Z/23/Z, 226052/Z/22/Z & 228186/Z/23/Z, the United Kingdom Research and Innovation (#APP8583), the Medical Research Foundation (MRF-RG-ICCH-2022-100069), UK International Development (301542-403), the Bill & Melinda Gates Foundation (INV-063472) and Novo Nordisk Foundation (NNF24OC0094346). The contents of this publication are the sole responsibility of the authors and do not necessarily reflect the views of the European Commission or the other funders. TCDL acknowledges funding from the National Institute for Health and Care Research (NIHR) Applied Research Collaboration East Midlands (ARC EM) and Leicester NIHR Biomedical Research Centre (BRC) and the Wellcome Trust (226080/Z/22/Z). The content and findings reported herein are the sole deductions, views, and responsibility of the researchers and do not necessarily reflect the official position and sentiments of the funding agencies.

## Supplementary Materials

**Table S1:** Brief summary of widely used approaches for arboviruses risk mapping.

| Arbovirus<br>Disease<br>Mapping Models | Input Data | Pros | Cons | Resolution | What questions it<br>answers | Public Health<br>Applications | Refs |
| --- | --- | --- | --- | --- | --- | --- | --- |
| Mechanistic<br>Models | Vector and virus lab<br>experiments data<br>(e.g vector lifespan,<br>biting rate,<br>incubation period,<br>human infectious<br>period) | Biologically informed;<br>mechanistic<br>understanding of<br>transmission | Do not incorporate<br>population dynamics<br>or immunity | Grid cell level<br>(climate<br>surfaces) | What are the<br>environmental limits<br>and optimal<br>conditions for<br>transmission? | Future projections<br>for mitigation and<br>adaptation | [26,59] |
| Vector occurrence<br>(ecological niche<br>models (ENMs)) | Vector Occurrence<br>points | Suitable for mapping<br>large areas and various<br>environmental<br>conditions, utilizing<br>geolocation of vector<br>occurrence data<br>combined with<br>high-resolution<br>environmental datasets | 1. Does not consider<br>transmission intensity<br>or pathogen presence<br>2. Outputs should be<br>used with caution due<br>to data gaps and<br>biases not reflected in<br>geographically precise<br>predictions | Fine spatial<br>resolution | Where are vector<br>species likely to<br>survive? | Target vector<br>surveillance and<br>control efforts | [5,55] |
| Disease<br>occurrence<br>(ENMs) | Dengue disease<br>occurrence points | Suitable for mapping<br>large areas and various<br>environmental<br>conditions, utilizing<br>geolocation of disease<br>occurrence data combined with<br>high-resolution<br>environmental datasets | 1. Tells us nothing<br>about magnitude of<br>risk of transmission<br>2. Outputs should be<br>used with caution due to data gaps and<br>biases not reflected in<br>geographically precise<br>predictions | Moderate to<br>high<br>resolution<br>depending<br>on resolution<br>of input data | Where can the viral<br>transmission of<br>dengue occur and<br>where might it<br>spread? | Early warning<br>systems; risk<br>communication | [6] |
| Reported disease<br>incidence<br>(Incidence-based<br>models) | Dengue case data | Grounded in<br>epidemiological data;<br>estimates intensity of<br>transmission - gives<br>best estimate of clinical<br>burden and treatment<br>resource needed | Biased by<br>underreporting;<br>missing data limits<br>accuracy, difficult to<br>compare across<br>countries due to<br>differing case<br>detection, diagnosis,<br>and reporting criteria | Administrativ<br>e units to<br>pixel level | 1. How does disease<br>burden vary across<br>space and time?<br>2. Which areas<br>should be prioritized<br>for intervention and<br>resource allocation? | Resource<br>allocation,<br>intervention<br>planning | [6,38,56] |
| Seroprevalence<br>and Cohort | Blood sample,<br>Demographic info,<br>Exposure history | Captures asymptomatic<br>& past infections;<br>enables temporal<br>analysis | Have a coarse<br>temporal resolution,<br>geographically sparse<br>and rare in<br>low-transmission<br>areas | Individual to<br>community<br>level | What proportion of<br>the population has<br>been infected? When<br>and where did<br>transmission occur? | Assess immunity<br>gaps; inform<br>vaccine strategy | [57,58] |

---

**Figure S1:**
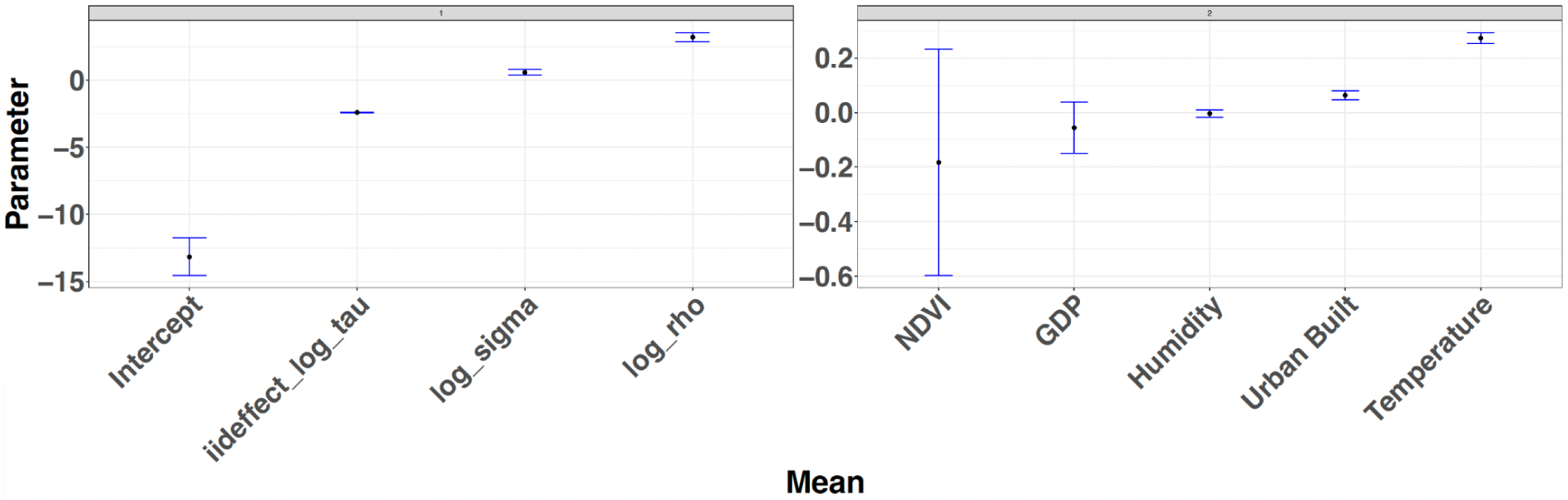
Posterior mean estimates and 95% credible intervals for model parameters from the disaggregation regression model. The left panel shows the intercept, spatial hyperparameters (log_rho, log_sigma), and the iid random effect (log_tau). The right panel displays fixed effects for the covariates used in the model.

**Figure S2:**
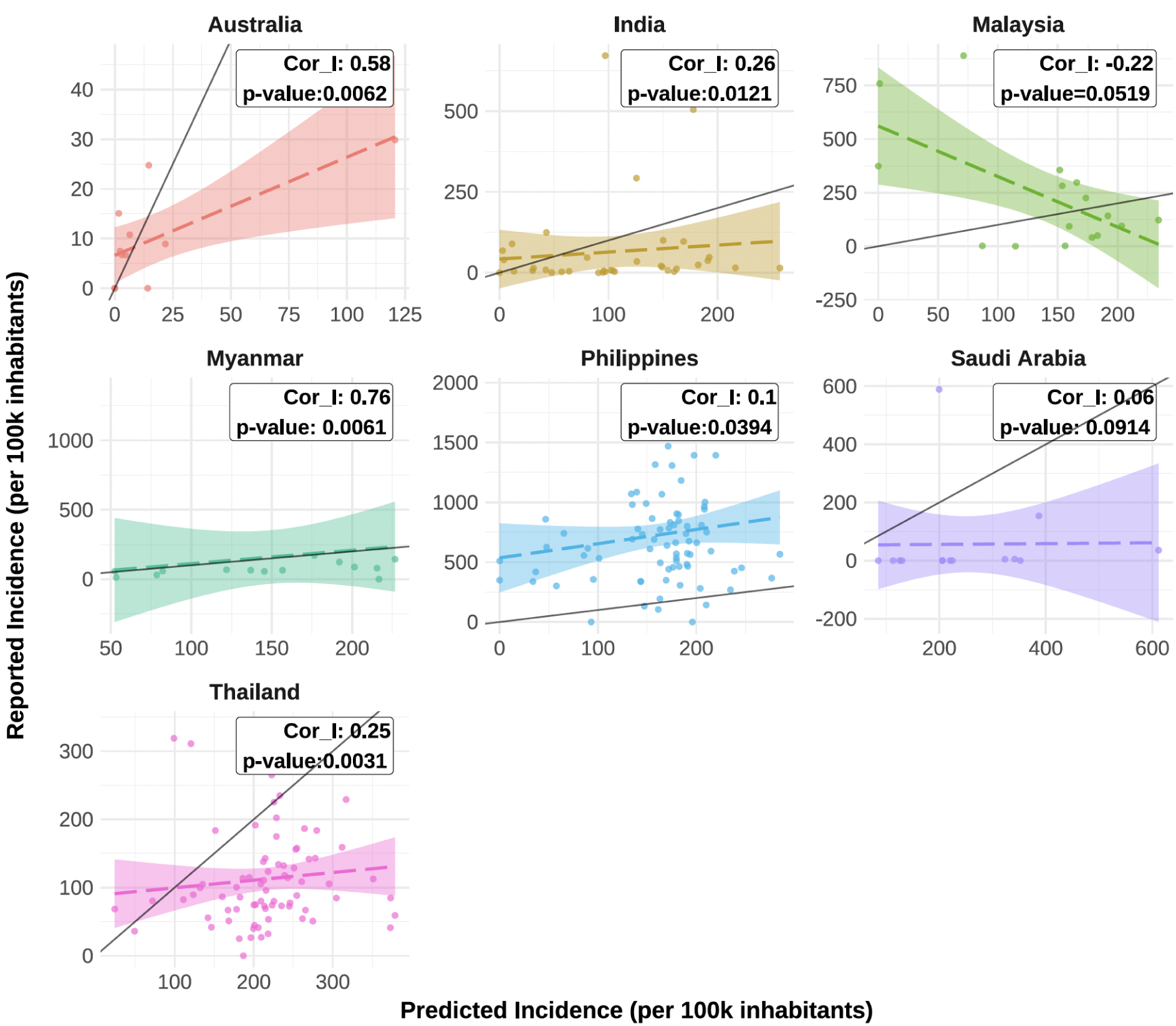
**Model performance across seven independent countries**. Scatter plots show the relationship between observed and predicted dengue incidence rates at the administrative level 1. For each country, we report Cor_I, the spearman correlation between observed and predicted incidence per 100k inhabitants with corresponding p-values. Shaded areas represent 95% confidence intervals around the regression line. Bottom panels show the posterior means and standard errors for selected covariate effects and spatial hyperparameters estimated from the disaggregation model.

**Figure S3:**
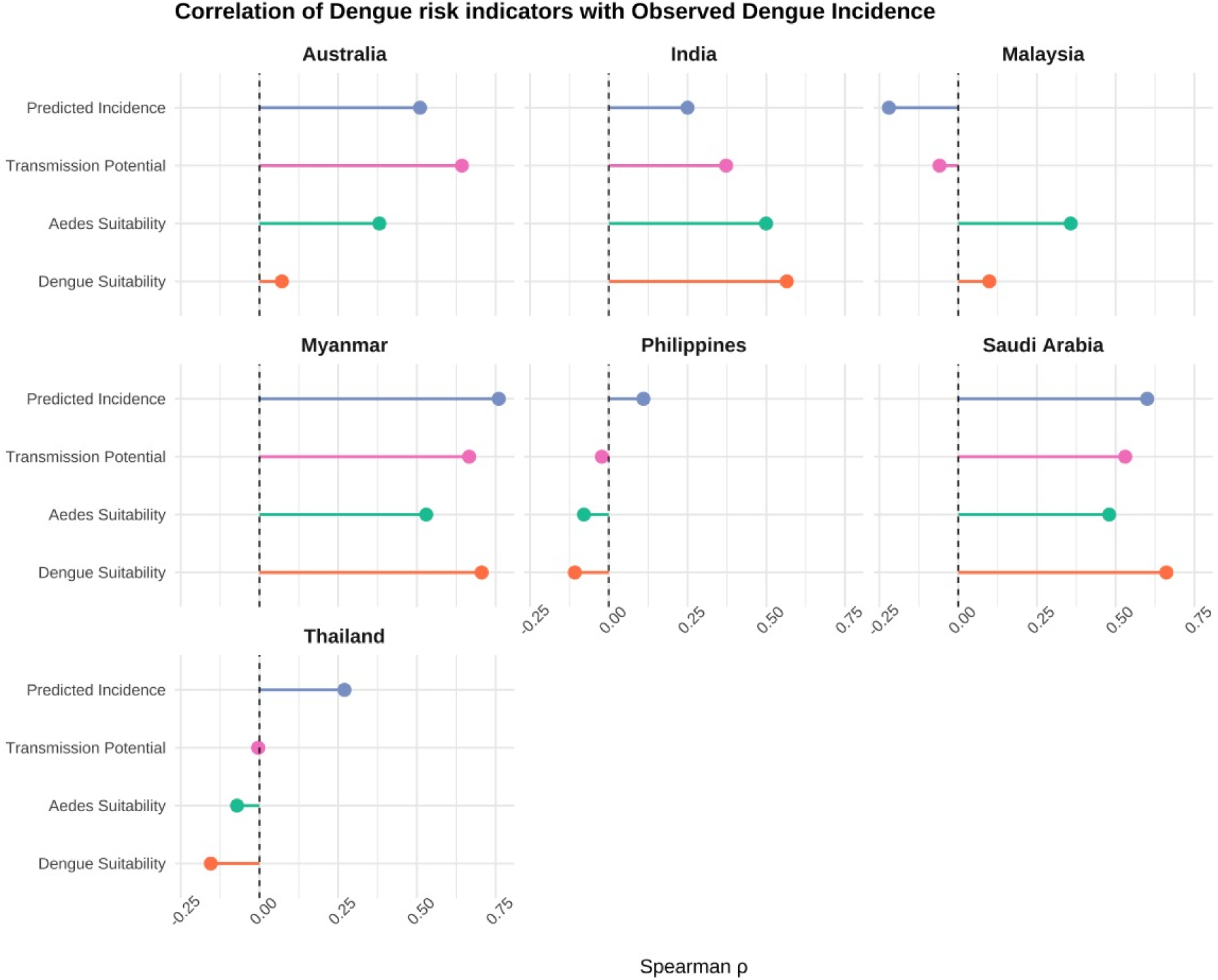
Spearman correlation (ρ) between observed dengue incidence and four risk indicators across seven countries. The plot shows correlations between nationally reported dengue incidence and (1) predicted incidence from our disaggregation regression model (blue), (2) transmission potential index (pink), (3) Aedes aegypti suitability index (green), and (4) dengue environmental suitability index (orange).

**Figure S4:**
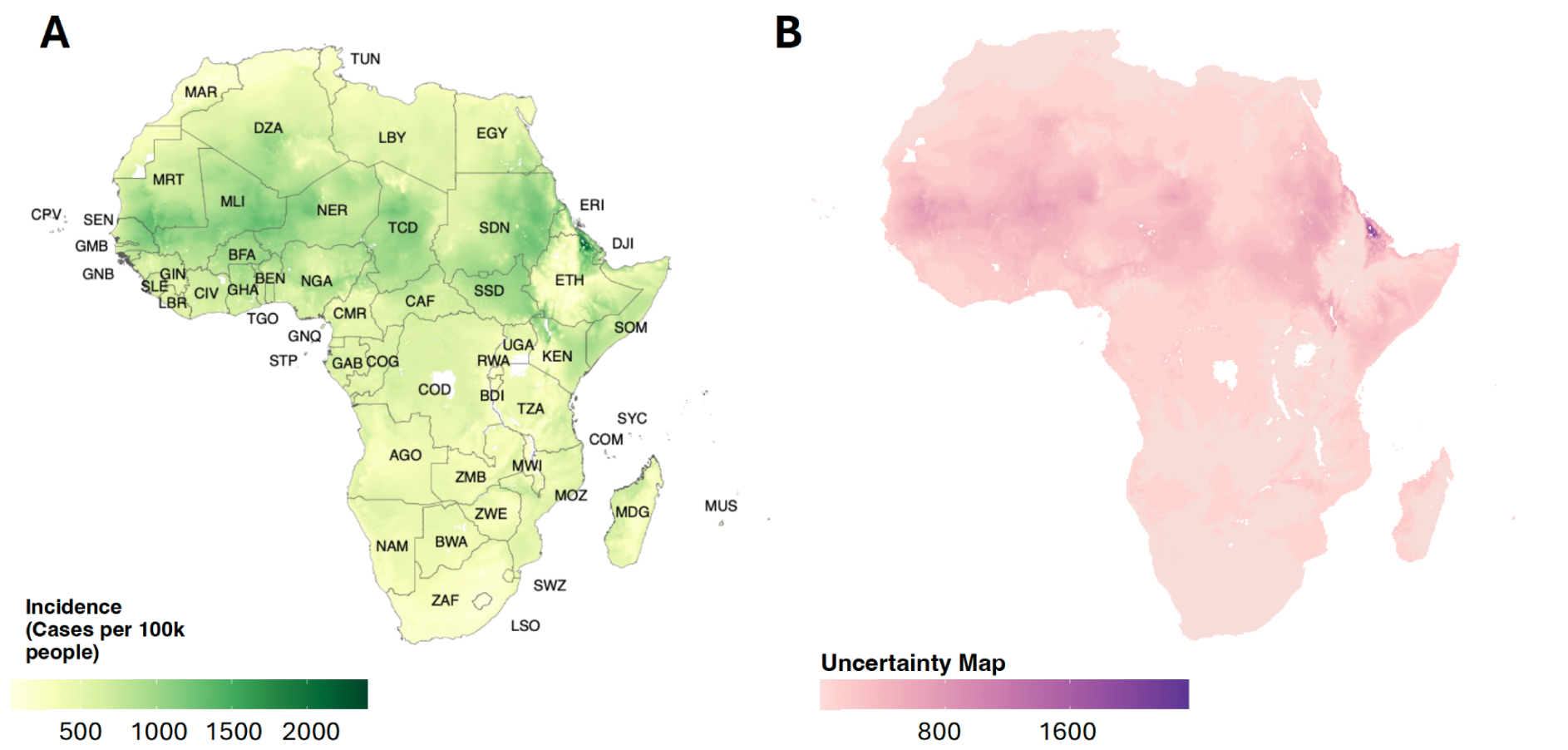
Panel (A) Spatial predictions of dengue incidence. (per 100k inhabitants) and uncertatinty plot across the African continent using the disaggregation regression model trained on case data from Central and South America. **Panel (B) shows the associated uncertainty,** calculated as the width of the 95% confidence interval (UCI – LCI). AGO = Angola, BEN = Benin, BFA = Burkina Faso, BWA = Botswana, CAF = Central African Republic, CIV = Côte d’Ivoire, CMR = Cameroon, COD = Democratic Republic of the Congo, COG = Republic of the Congo, COM = Comoros, CPV = Cabo Verde, DJI = Djibouti, DZA = Algeria, EGY = Egypt, ERI = Eritrea, ETH = Ethiopia, GAB = Gabon, GHA = Ghana, GIN = Guinea, GMB = Gambia, GNB = Guinea-Bissau, GNQ = Equatorial Guinea, KEN = Kenya, LBR = Liberia, LBY = Libya, LSO = Lesotho, MDG = Madagascar, MLI = Mali, MOZ = Mozambique, MRT = Mauritania, MUS = Mauritius, MWI = Malawi, NAM = Namibia, NER = Niger, NGA = Nigeria, RWA = Rwanda, SDN = Sudan, SEN = Senegal, SLE = Sierra Leone, SOM = Somalia, SSD = South Sudan, STP = São Tomé and Príncipe, SWZ = Eswatini, SYC = Seychelles, TCD = Chad, TGO = Togo, TUN = Tunisia, TZA = Tanzania, UGA = Uganda, ZAF = South Africa, ZMB = Zambia, ZWE = Zimbabwe.

**Figure S5:**
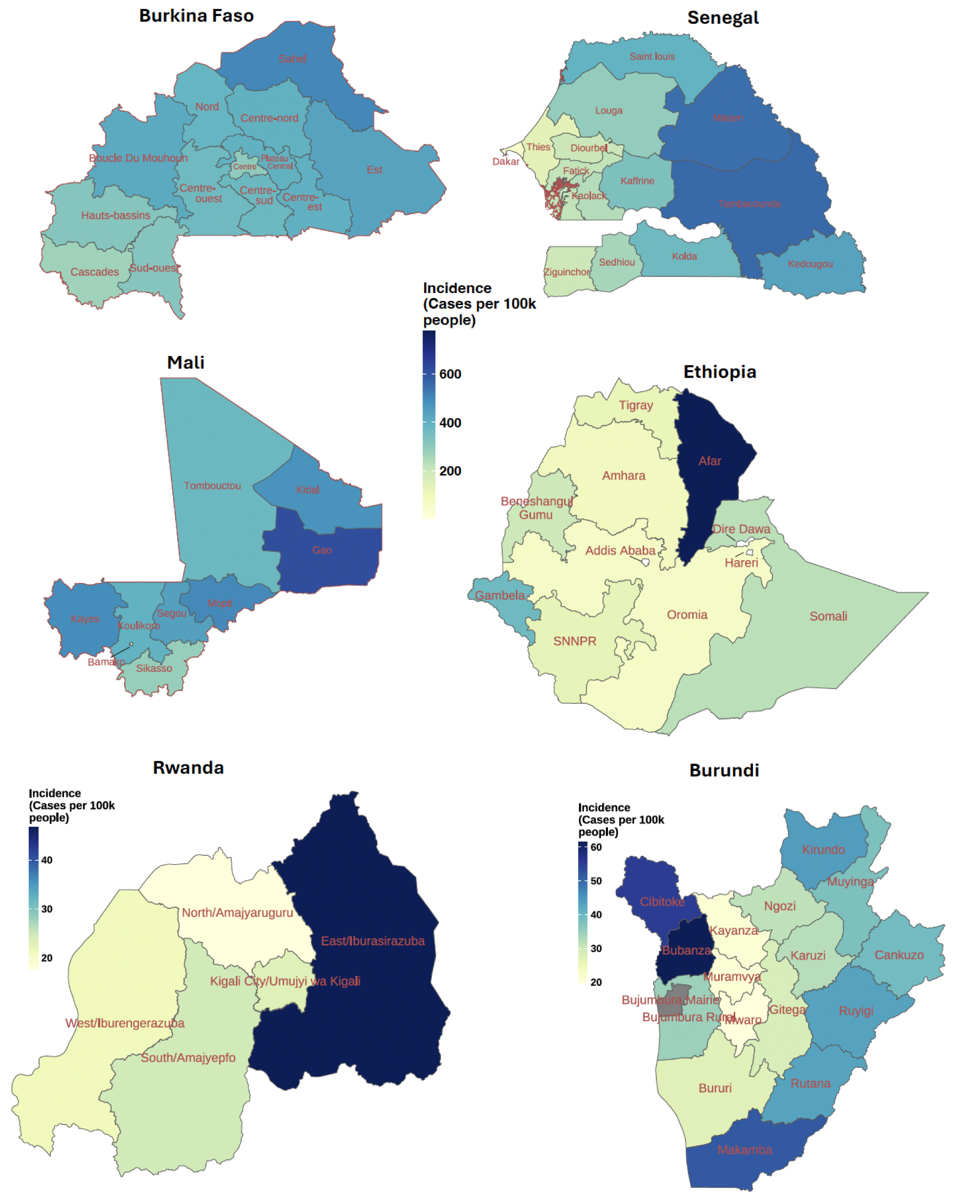
**Subnational predicted dengue incidence in selected countries in Africa**. Maps show predicted cases per 100,000 people at the administrative level 1 for Burkina Faso, Senegal, Mali, Ethiopia, Rwanda and Burundi.

**Figure S6:**
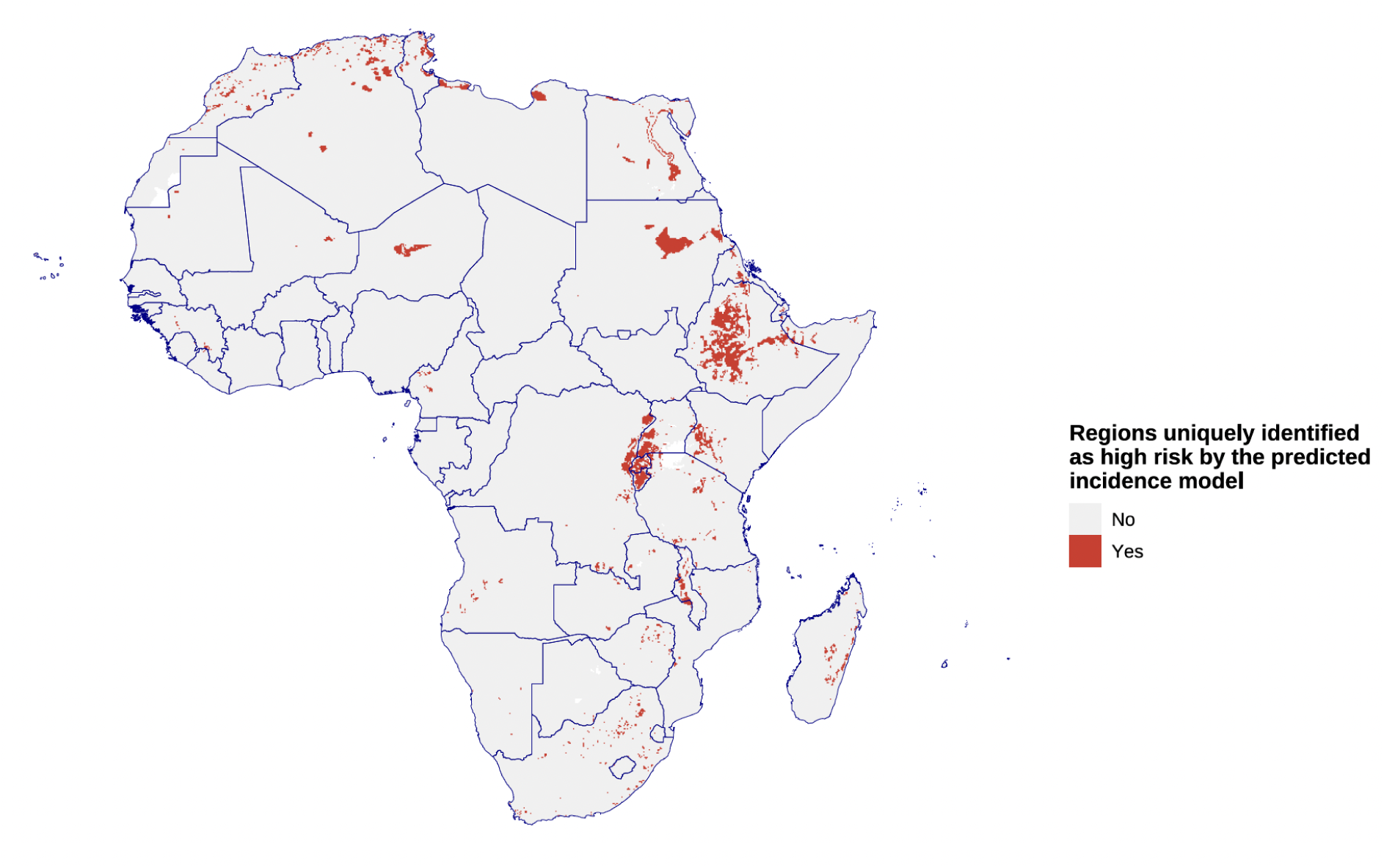
Spatial distribution of regions uniquely identified as high risk by the incidence based model. Areas shown in red indicate pixels where only the disaggregation-derived incidence model predicted high dengue risk (after accounting for the threshold).

**Figure S7:**
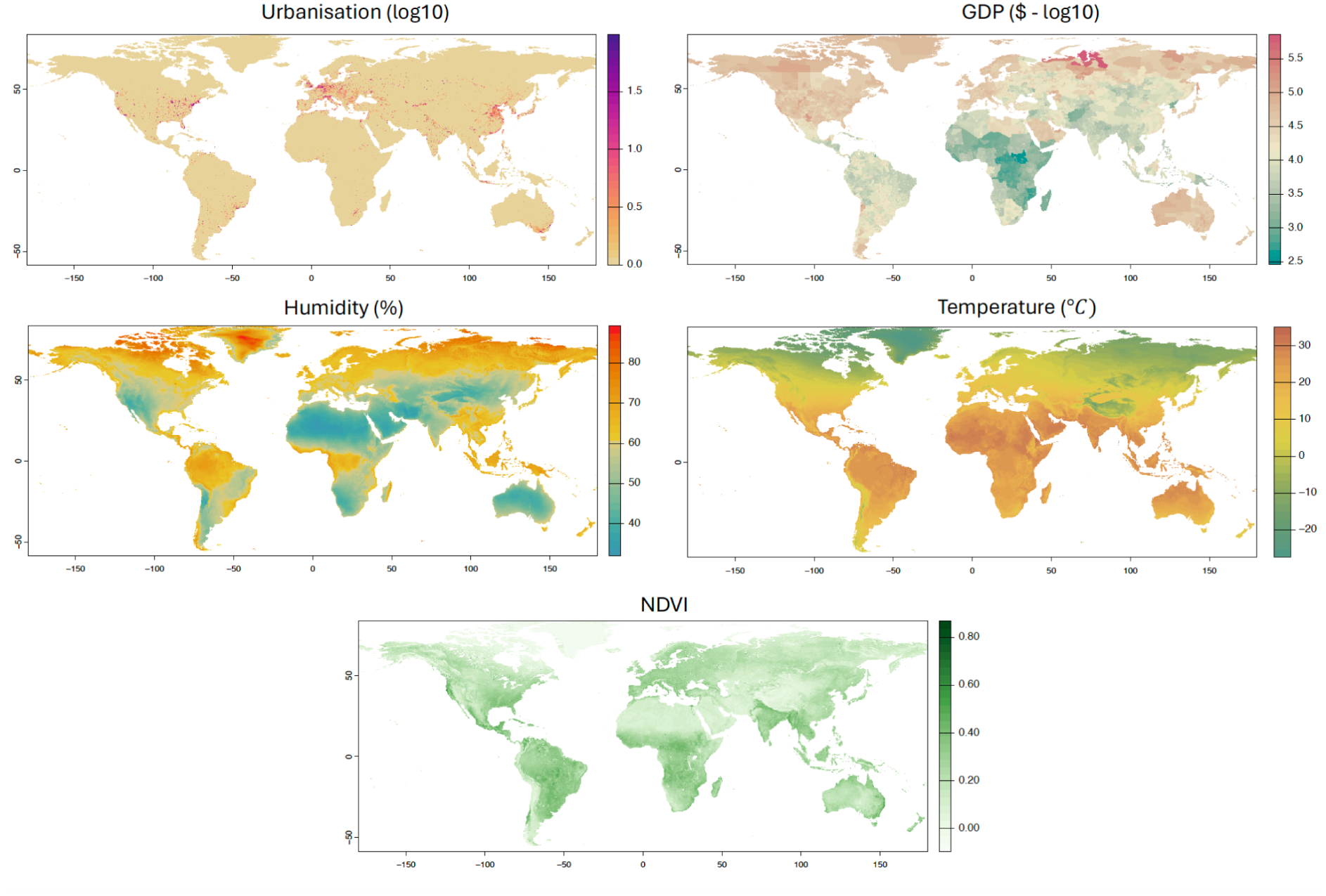
Maps of covariates used in the study: Urbanisation, GDP, Humidity, Temperature and Normalized Difference Vegetation Index (NDVI).

